# The true size of placebo analgesia: Concordant neural and behavioural measures of placebo analgesia during experimental acute pain

**DOI:** 10.1101/412296

**Authors:** E Valentini, SM Aglioti, B Chakrabarti

**Affiliations:** Department of Psychology and Centre for Brain Science, University of Essex, UK; Sapienza Università di Roma, Dipartimento di Psicologia, Italy; Fondazione Santa Lucia, Istituto di Ricovero e Cura a Carattere Scientifico, Italy; Centre for Integrative Neuroscience and Neurodynamics, School of Psychology and Clinical Language Sciences, University of Reading, UK.

**Keywords:** Electroencephalography, laser evoked potentials, lidocaine, nociception, pain, placebo analgesia, vaseline

## Abstract

‘Placebo analgesia’ refers to the reduction of pain following the administration of an inactive treatment. While most clinical trials compare a drug treatment against a placebo to determine the efficacy of the analgesic, most experimental studies of placebo analgesia do not include a real analgesic condition. A direct comparison of placebo against a real analgesic can inform us about the true size of the placebo effect. To this end, we aimed to provide a robust estimate of placebo analgesia by contrasting the effect of pain relief expectation from an inert cream (vaseline) against a real topical analgesic agent (lidocaine) applied on two different limbs and their respective control conditions. Pain reports and electroencephalography (EEG) responses triggered by laser nociceptive stimulation were collected. Forty typical healthy adults were enrolled in a double-blind randomized within-subject study where a standard placebo induction script of verbal suggestions in a sham medical setting was used to enhance the expectation on treatment outcome. In line with the earliest studies of placebo analgesia, majority (30 of 40) of participants was placebo responders, i.e. they reported lower pain to the placebo treatment. Placebo responders reported low pain and displayed low laser evoked potentials (LEPs) amplitude for both the analgesic and placebo treatment limbs compared to the respective control limbs. Placebo analgesia correlated positively with the amplitude of the LEPs, thus establishing convergent validity of the findings. This study provides a robust estimate of the neural and behavioural measures of placebo analgesia, in comparison to a real analgesic. These estimates can help inform the quantitative criteria for similar neural and behavioural measures in assessing the effectiveness of a real drug in placebo controlled trials.

## Introduction

Placebo effects lead an individual to display/feel an experiential improvement following the administration of an inert treatment with no actual therapeutic properties. In other words, factors differing from the purported treatment can cause a beneficial physical response. This has been observed in several clinical conditions and diseases, particularly in clinical pain (Tuttle et al., 2015). While the phenomenon is well recognized, the magnitude of placebo effects, the influence of the context, and their temporal course are less known (see Benedetti, 2008 for a general review).

Despite a robust body of evidence over the last four decades starting from (Levine, Gordon, & Fields, 1978), there remain important concerns on the robustness and reliability of placebo, especially in the clinical settings. Meta-analytic studies have indicated the presence of potential confounds (e.g. regression to the mean; Artus, van der Windt, Jordan, & Hay, 2010; Hrobjartsson, Kaptchuk, & Miller, 2011) that led to overestimation of very small to null placebo effects (Hrobjartsson & Gotzsche, 2001, 2004, 2010; Hrobjartsson et al., 2011). Notwithstanding considerable individual variability in the magnitude of placebo analgesia (Wager, Atlas, Leotti, & Rilling, 2011), several studies indicate that placebo analgesia is a reliable and consistent phenomenon (Atlas & Wager, 2014; Finniss, Kaptchuk, Miller, & Benedetti, 2010; Price et al., 1999; Vase et al., 2015). Interestingly, clinical trials for analgesics and experimental studies of placebo pose a methodological contrast. While clinical trials for analgesics routinely compare them against a placebo to estimate the magnitude of the analgesic effect, most experimental studies of placebo analgesia do not use a real analgesic treatment to estimate the size of the placebo effect (e.g. Price et al., 1999, but see Vase, Robinson, Verne, & Price, 2005 for an exception). Here we address this methodological difference by directly comparing the magnitude of placebo analgesia against that of a known analgesic.

Laser thermal stimulation provides a targeted way to selectively stimulate nociceptive free nerve endings in the skin. In particular, solid state lasers (as the one used in the current study) offers a reduced risk of superficial burns than the CO_2_ laser, due to its shorter wavelength (1.34 μm). In addition, solid state lasers allow a better afferent-volley synchronization which results in enhanced amplitudes and shorter latencies of cortical responses (Perchet et al., 2008). To date, recording of electroencephalographic activity during laser thermal stimulation (Laser Evoked Potentials, LEP) provides the most reliable and selective neurophysiological method of assessing the function of nociceptive pathways (Garcia-Larrea, 2012). However, there is still relatively little research using laser thermal stimulation to study placebo analgesia.

Using LEP, here we aimed to provide a robust estimation of placebo analgesia by contrasting the effect of pain relief expectation from an inert cream (vaseline) against a real topical analgesic agent (lidocaine) and their respective control conditions in a large sample of healthy volunteers (n=40). We collected pain reports and EEG responses triggered by laser nociceptive stimulation in a double-blind randomized within-subject design whereby healthy volunteers underwent a standard placebo induction script of verbal suggestions in a sham medical setting meant to enhance the expectation on treatment outcome. Verbal induction of expectations about the outcome can not only lead to formation of conscious expectations, but also bring online effects of unconscious learning, two processes that can lead to placebo analgesia (e.g. Benedetti et al., 2003; Pecina, Stohler, & Zubieta, 2014).

## 2. Material and Methods

### 2.1 Subjects

EEG data were collected from 40 healthy volunteers. We excluded one participant from data analysis as she questioned about covert experimental aims possibly concerning the investigation of placebo in the debriefing phase. The remaining 39 participants (21 females) were aged 24.9±4.5 (mean±SD). All had normal or corrected-to-normal vision and were naïve as to the purpose of the experiment. None of the participants had a history of neurological or psychiatric illnesses or conditions that could potentially interfere with pain sensitivity (e.g. drug intake or skin diseases). Participants gave written informed consent and were debriefed about the actual aim of the study at the end of the experiment. The participants could therefore decide to withdraw their consent about data usage if they wished so. All experimental procedures were approved by the Fondazione Santa Lucia ethics committee and were in accordance with the standards of the Declaration of Helsinki. No participant had short or medium term symptoms (e.g. Inflammation) associated with the compounds used in this study.

### 2.2 Nociceptive stimulation

Radiant-heat stimuli were generated by an infrared neodymium yttrium aluminium perovskite (Nd:YAP) laser with a wavelength of 1.34 μm (Electronical Engineering, ElEn, Florence, Italy). Laser pulses selectively and directly activate the Aδ and C-fiber nociceptive terminals located in the superficial layers of the skin (Cruccu et al., 2003). Laser pulses were directed at the dorsum of both left and right hand and foot, on a squared area (5×5 cm) defined prior to the beginning of the experimental session and highlighted using a He-Ne guide laser. The laser pulse (3 ms duration) was transmitted via an optic fibre and its diameter was set at approximately 5 mm (28 mm^2^) by focusing lenses. After each stimulus, the laser beam target was shifted by approximately 1 cm in a random direction, to avoid nociceptor fatigue or sensitization.

Before the recording session, a familiarization and calibration procedure was carried out to check the quality of the sensation associated with radiant heat stimuli. In this procedure, the energy of the laser stimulus was individually adjusted using the method of limits (laser step size: 0.25 J), separately for each of the four stimulated territories (left hand, right hand, left foot, right foot). During this procedure subjects were asked to report the quality and the intensity of the sensation elicited by each laser pulse using a numerical rating scale (NRS, ranging from 0=no sensation, to 8=unbearable pain). The energy of laser stimulation needed to achieve a rating of 6 (corresponding to ‘moderate pain’) was chosen as experimental energy value. We checked that this value corresponded to a rating of about 60 on visual analogue scale (VAS) ranging from 0 (not painful) to 100 (extremely painful). Once nociceptive intensity was calibrated, participants underwent a brief familiarization block of 10 stimuli. Importantly, there was no difference in the average energy used to obtain a moderate sensation of pain for both feet and hands: right and left hand, 2.27±0.34 J; right and left foot, 2.33±0.32 J. According to the parameters mentioned above, laser pulses elicited a clear pinprick/burning brief sensation of acute pain related to the activation of Aδ and C fibres.

### 2.3 EEG recording

The electroencephalogram (EEG) was recorded using 54 tin scalp electrodes placed according to the International 10-20 system, referenced against the nose and grounded at AFz. Electro-oculographic (EOG) signals were simultaneously recorded using surface electrodes. Electrode impedance was kept below 5 KΩ. The EEG signal was amplified and digitized at a sampling rate of 1,000 Hz.

### 2.4 Experimental design

Upon arrival participants were welcomed in a temperature-controlled room by two experimenters (EV, BC) dressed in white coats. They introduced the participants to the study using the same set of sentences (see Appendix), and informed them about the whole procedure. In brief, participants were told that two analgesics (named *Varicaine* and *Exacaine*) were being evaluated for their efficacy. In reality, one of these was an inert cream (vaseline, labelled as cream A and called *Varicaine*), while the other was a topical analgesic (5% lidocaine, labelled as cream B and called *Exacaine*).

Participants then underwent the EEG cap montage. The analgesic cream was applied on the dorsal surface of one of four limbs (hand/foot, coded as Treat B). An identical site in the contralateral limb was used as its control (no cream, control site, coded as Ctrl B). Same procedure was adopted for the inert cream on the other pair of limbs (coded as Treat A and Ctrl A respectively).

The conditions were counterbalanced in a double-blind fashion across participants (Fig. 1).

**Fig 1.**
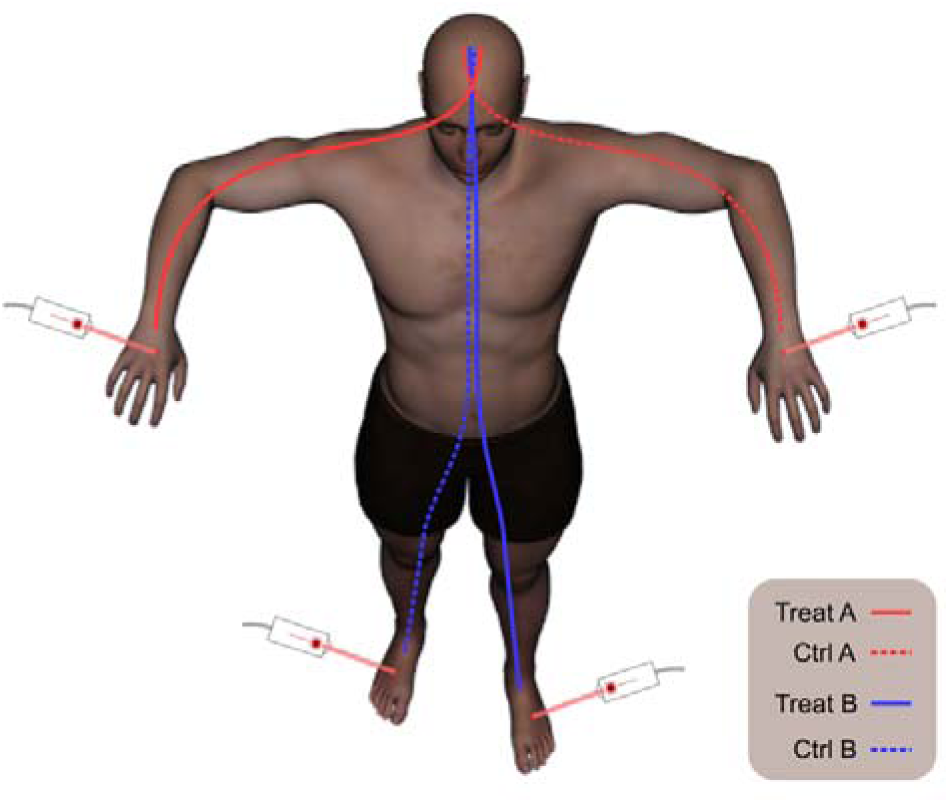
Participants were told that two analgesics were being evaluated for their efficacy. They were unaware that one of these was an inert cream (vaseline, labelled as cream A and called Varicaine), while the other was an actual topical analgesic (lidocaine, labelled as cream B and called Exacaine). Subjective pain thresholds for moderate pain (a rating of 6 out of 10) was established for each participant before the application of creams. The analgesic cream was applied on the dorsal surface of one of four limbs (hand/foot, coded here as Treat B). An identical site in the contralateral limb was used as its control (no cream, control site, coded here as Ctrl B). The same procedure was adopted for the inert cream on the other pair of limbs (coded here as Treat A and Ctrl A respectively). The conditions were counterbalanced in a double-blind fashion across participants. Each block lasted between 10 and 15 min, and an interval of 5 min separated the two blocks. In each block we delivered 30 laser pulses, using an inter-stimulus interval ranging between 5 and 15 s. At the end of each train of 10 stimuli, participants were asked to rate the intensity of the painful sensation elicited by the laser stimuli using a visual analogue scale ranging from 0 (not painful) to 100 (extremely painful).

Creams were spread and left acting on the skin for a mean duration of 13:52 min (SD=2.52 min). After careful rubbing of the creams off the administration sites, all four limbs were stimulated using a Nd:YAP laser at an energy level corresponding to subjective threshold for moderate pain (i.e. NRS=6). Participants were asked to focus their attention on the painful stimuli while closing their eyes and relax their muscles. Laser-evoked EEG responses were obtained following the stimulation of the dorsum of the right and left hand and foot in four separate blocks, on the same day. Each block lasted between 10 and 15 min, and an interval of 5 min separated the two blocks. In each block we delivered 30 laser pulses, using an inter-stimulus interval (ISI) ranging between 5 and 15s. At the end of each train of 10 stimuli, participants were asked to rate the pain intensity and unpleasantness of the painful sensation elicited by the laser stimuli using a visual analogue scale (VAS) ranging from 0 (no sensation, no unpleasant at all) to 100 (intolerable intensity/intolerable unpleasantness).

At the end of the experiment, participants went through a structured debriefing interview in which we asked their opinion on the experimental aims (e.g. “What do you think was the study objective?” and “Did you notice any difference in the efficacy of the two creams?”) and were debriefed regarding the deception.

### 2.5 Data analysis

#### 2.5.1 General statistical approach

Dependent variables were analyzed with repeated-measures Analysis of Variance (ANOVA) with factors ‘expectation’ (treatment, no treatment) and treatment ‘label’ (A – placebo, B – analgesic). Further, we run an additional ANOVA only on placebo responders, i.e. individuals who reported significant lower pain unpleasantness during placebo vs. no treatment (n= 30). The choice of pain unpleasantness as the variable of interest was supported by the evidence that the major feature of the multidimensional pain experience is its affective quality rather than its intensity (Merskey, Bogduk, & Pain, 1994).

Statistical analyses were performed using Statistica^®^ 8.0 (StatSoft Inc., Tulsa, Oklahoma, USA). Variability is reported as standard error of mean (SEM) unless reported otherwise. The level of significance was set at *p*<0.05. We reported Cohen’s d and partial eta squared (pη^2^) as measures of effect size. Tukey HSD tests were used to perform post-hoc pairwise comparisons.

#### 2.5.2 Laser evoked potentials

EEG data were processed with EEGLAB (v.12; Delorme & Makeig, 2004 and Letswave 5, http://nocions.webnode.com/). Single participant data were merged in a unique experimental session file and down-sampled to 250 Hz. Sinusoidal artifacts (50-100 Hz) were then removed using CleanLine, an EEGLAB plugin which enabled us to selectively delete power line frequency contribution from the recorded signal (http://www.nitrc.org/projects/cleanline). Further, signal was DC removed and band-pass filtered from 1 to 30 Hz (filter order: 4). Data were then segmented into epochs using a time window ranging from 1 s before to 2 s after the stimulus (total epoch duration: 3 s) and baseline corrected using the mean of the entire epoch (Groppe, Urbach, & Kutas, 2011). Epoched data were merged and further processed using independent component analysis (ICA; Vigário, 1997) to subtract EOG and muscle-related artifacts, aided by the semi-automatic approach offered by Adjust (Mognon, Jovicich, Bruzzone, & Buiatti, 2011), an EEGLAB plugin which identifies artifactual independent components using an automatic algorithm that combines stereotyped artifact-specific spatial and temporal features. After ICA and an additional baseline correction (−500 to 0 ms), we re-referenced data to a common average reference (Lehmann & Skrandies, 1980) and segmented in four average waveforms time-locked to the stimulus onset, one for each experimental condition (Ctrl A; Treat A; Ctrl B; Treat B). Single-subject average waveforms were subsequently averaged to obtain group-level average waveforms. Group-level scalp topographies were computed by spline interpolation. Scalp topographies were plotted at the peak latency of the N2 and P2 LEP waves, measured at the vertex (Cz electrode). The N2 wave was defined as the most negative deflection after stimulus onset. The P2 wave was defined as the most positive deflection after stimulus onset. We used group-level median peak values to identify the temporal window to extract the minimum (N2, 180-280 ms) and the maximum (P2, 280-480 ms) amplitudes for each participant. These two waves seem to result from sources in bilateral operculo-insular and anterior cingulate cortices (Garcia-Larrea, Frot, & Valeriani, 2003). They are significantly modulated by both top-down and bottom-up attentional factors (reviewed in Legrain et al., 2012).

#### 2.5.3 Correlation between pain ratings and N2-P2 amplitudes

Placebo and analgesia response magnitude was calculated as a ratio of the average ratings, N2-P2 peak-to-peak amplitude, for the placebo control limb divided by that for the placebo treatment limb (Ctrl/Treat). In other words, the greater the value of this ratio the greater the analgesic effect.

## 3 Results

### 3.1 Psychophysics

All participants described the sensation elicited by the laser stimuli as clearly painful and pricking. The average ratings (mean±SD) of the pain unpleasantness for each experimental condition as well as the effect sizes are reported in Table 1.

**Table 1.** Mean (±SD) of pain ratings (unpleasantness)(top) in the full sample. Cohen’s d as for both types of ratings as well as for the ratio Ctrl/Treat (bottom). A refers to the inert cream, and B refers to the real analgesic. Ctrl A refers to the no-treatment contralateral limb control for the inert cream; Ctrl B refers to the no-treatment contralateral limb control for the real analgesic.

|  | Pain rating (unpleasantness) |  |  |  |
| --- | --- | --- | --- | --- |
|  | Treat A | Ctrl A | Treat B | Ctrl B |
| **RATINGS** | 62.27<br>(±18.14) | 70.27<br>(±15.76) | 56.81<br>(±17.73) | 60.94<br>(±18.13) |
|  | Pain rating (unpleasantness) |  |  |  |
|  | Treat A<br>vs. Ctrl<br>A | Treat B<br>vs. Ctrl B | Ctrl A/Treat A vs.<br>Ctrl B/Treat B |  |
| **EFFECT**<br>**SIZE** | -0.51 | -0.21 | -0.07 |  |

#### 3.1.2 Effects of expectation and treatment label

The ANOVA performed on the unpleasantness ratings revealed main effects of both ‘expectation’ (*F*_38_=19.62; *P*<0.001; pη^2^=0.34) and treatment ‘label’ (*F*_38_=6.70; *P*=0.01; pη^2^=0.15), but no significant interaction between the two factors (*F*_38_=2.21; *P*=0.14; pη^2^=0.05). This pattern of results indicates that participants felt less pain unpleasantness when expecting treatment compared to no treatment and felt less pain unpleasantness during the analgesic-related (Ctrl B and Treat B) vs. placebo-related (Ctrl A and Treat A) stimulation (Fig. 2, A). The analysis on responders (Fig. 2, B) revealed no main effect of this ‘label’, suggesting that individuals responding better to the placebo treatment had no different unpleasantness depending on the type of cream used and its related control stimulation (*F*_29_=2.40; *P*=0.13; pη^2^=0.08) but rather showed lower pain unpleasantness when treatment was expected (*F*_29_=36.80; *P*<0.001; pη^2^=0.56) and with both ‘expectation’ and ‘treatment label’ (*F*_29_=7.83; *P*=0.009; pη^2^=0.21). These interactions reflect (i) a larger reduction of pain unpleasantness in responders when expecting the Treat A (i.e. *Varicaine*) compared to Ctrl A (58.56 vs. 71.21; *P*<0.001), (ii) a greater pain unpleasantness in responders during the Ctrl A against Treat B and Ctrl B (71.21 vs. 57.51 and 61.71; *Ps*<0.001).

**Fig 2.**
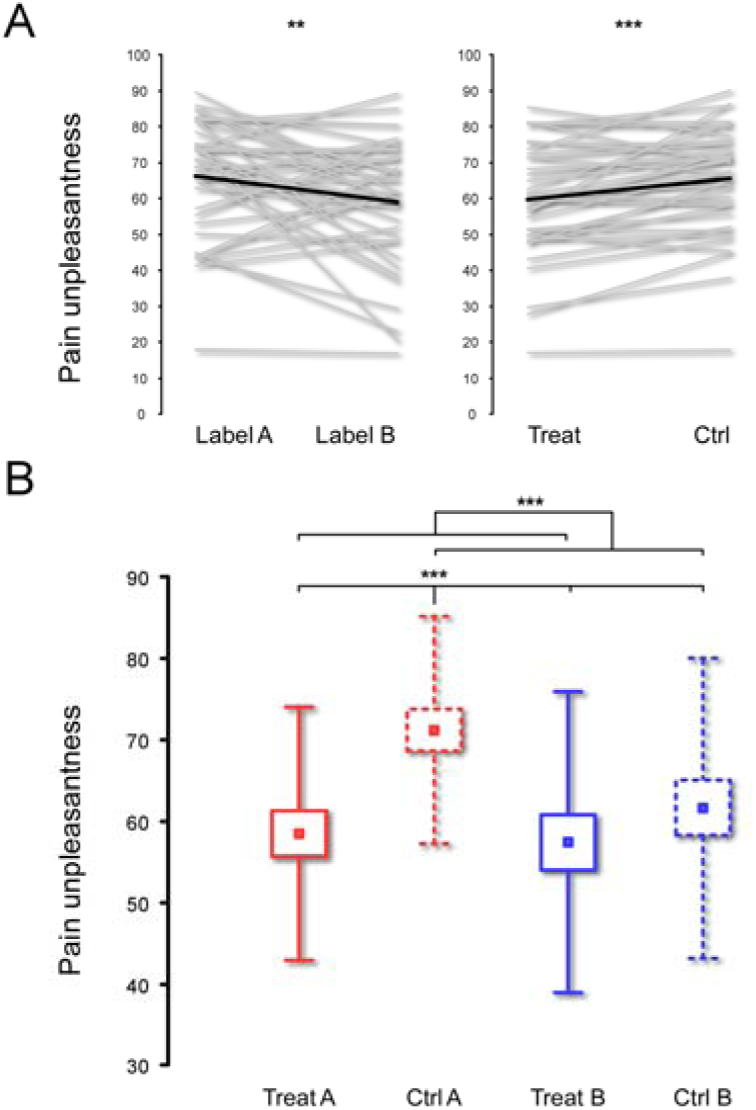
Panel A shows single subject average ratings of pain unpleasantness for the two levels (Label A, Label B) of factor treatment “label” (left) and the two levels (Treat, Ctrl) of the factor treatment “expectation” (right). Grand-average is shown with bold black line. Individuals reported lower pain unpleasantness during both placebo and analgesia treatment than in the respective control conditions (***p<0.001). They also reported lower pain unpleasantness during both actual analgesia and its control condition than during placebo and its control condition (***p≤0.001). Panel B shows results only for placebo responders. Box-plots show (mean ±SE±SD) of pain intensity ratings. The pattern observed in the full sample was enhanced in this subgroup (***p<0.01).

### 3.2 Laser evoked potentials

Fig. 3 (A) displays the grand average waveforms and global field power (GFP) of LEPs. Nociceptive stimuli delivered in the four conditions elicited maximal N2 and P2 waves at the electrode Cz with topographies maximally expressed over the scalp vertex (Fig. 3, A, top).

**Fig 3.**
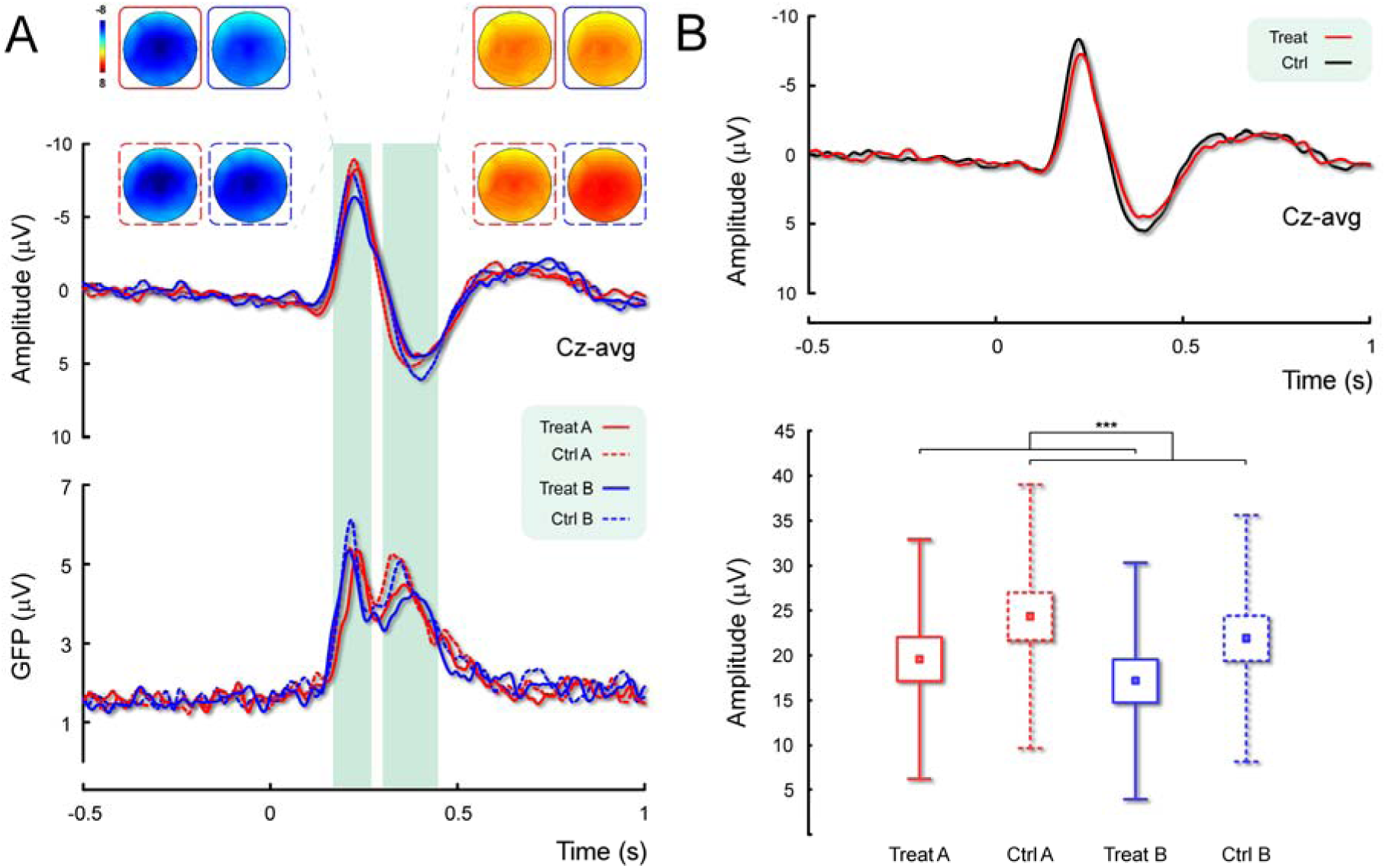
Panel A shows group-level average LEPs and scalp topographies of peak amplitudes (top) within the N2 and P2 latency range (180-280 and 280-480 ms post-stimulus respectively) as well as global field power (GFP; bottom) in the four conditions (Placebo-related in red, analgesia-related in blue; treatment in solid and control conditions in dashed lines). Note the greater amplitudes elicited by the stimulation of the no-treatment (control) limbs. Panel B clarifies this pattern by showing the main effect of treatment expectation on the vertex LEPs in the full sample (top). Box-plots (mean ±SE±SD) show N2-P2 peak-to-peak amplitude in placebo responders in the four conditions (bottom). Note the amplitude reduction in Treat A and B compared to Ctrl A and Ctrl B respectively.

#### 3.2.1 Effects of expectation and treatment label on N2-P2

The ANOVA performed on the peak-to-peak amplitude of the main vertex potentials N2-P2 extracted at the Cz electrode revealed a main effect of ‘expectation’ (*F*_38_=11.54; *P*=0.002; pη^2^=0.23) but no effect of treatment ‘label’ (*F*_38_=0.69; *P*=0.41; pη^2^=0.02) or interaction between the two factors (*F*_38_=0.98; *P*=0.33; pη^2^=0.02). This pattern of results indicates that participants displayed lower vertex potentials amplitude when expecting treatment compared to no treatment (Fig. 3, B). Peak-to-peak amplitudes in responders (Fig. 3, B, bottom) revealed no main effect of treatment ‘label’, suggesting that individuals responding better to the placebo treatment had no different N2-P2 LEP amplitude depending on the type of cream used and its related control stimulation (*F*_29_=0.79; *P*=0.38; pη^2^=0.03) but rather showed lower N2-P2 amplitude when treatment was expected (*F*_29_=24.26; *P*<0.001; pη^2^=0.45). However, there was no interaction between the two factors (*F*_29_<0.001; *P*=0.99; pη^2^<0.001).

### 3.3 Correlation of pain ratings with LEPs

The magnitude of the placebo response was calculated as the ratio of unpleasantness ratings of the control limb divided by that of the treatment limb (Fig. 4). This magnitude was positively correlated with the N2-P2 response, calculated similarly (i.e. N2-P2 response of the control limb divided by that of the treatment limb) (r_38_=0.50; *P*=0.001).

**Fig 4.**
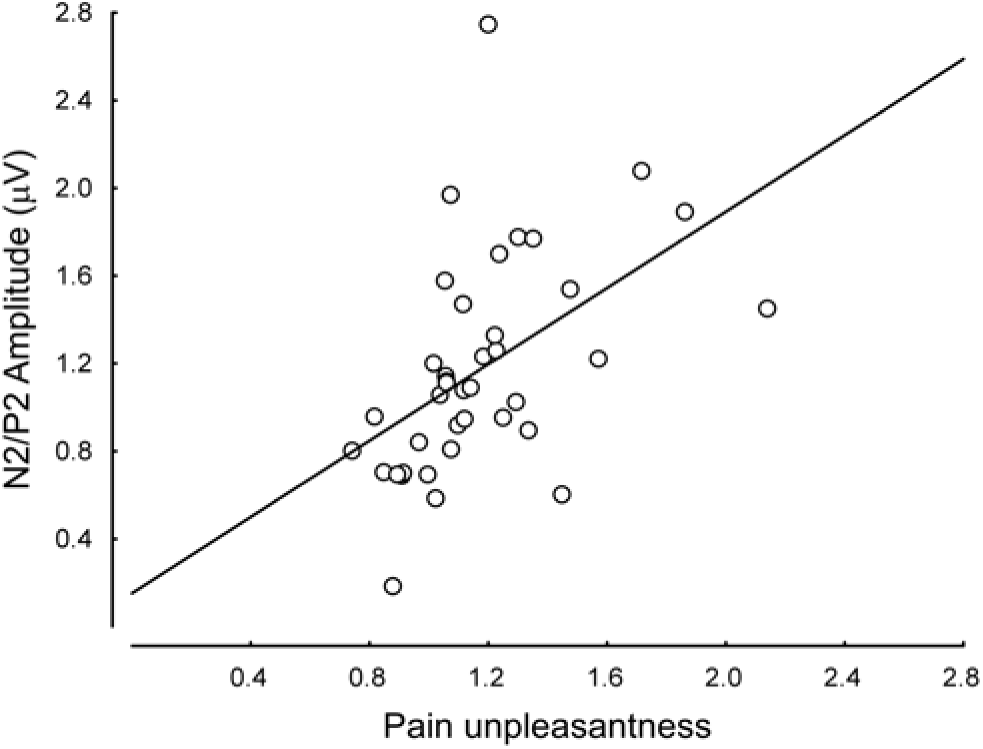
Scatterplot representing the relationship between ratings of pain unpleasantness (x-axis) and the amplitude of the N2/P2 LEPs (y-axis) with its linear fit. Both measures were calculated as a ratio of the control limb divided by treatment limb, thus providing an index of the placebo effect. Note that the reduction of pain unpleasantness associated with placebo is linked to a decrease of N2/P2 peak-to-peak amplitude.

## 4. Discussion

In this study, we estimated the magnitude of placebo analgesia against a real analgesic, using self-report and laser-evoked potential measures. Our results show that healthy volunteers felt less pain and displayed lower magnitude of EEG responses when receiving a purported analgesic treatment (regardless of whether this was a sham or actual analgesic compound) compared to the stimulation of non-treated skin territory (Figs 2 and 3). Magnitude of the placebo response computed from pain unpleasantness ratings was positively correlated with that computed from the neural response to placebo (Fig. 4). This study presents one of the few concordant behavioural and neural estimates of the placebo analgesia effect, using a true analgesic and a sham treatment and expand current knowledge about placebo analgesia and its neural correlates (Geuter, Koban, & Wager, 2017; Wager & Atlas, 2015 for reviews). Despite the high number of placebo responders (n=30 according to our identification criterion), the true effect size for the placebo effect was small (Table 1, d=−0.07). This is consistent with previous research (Price, Finniss, & Benedetti, 2008). Across all participants, our data demonstrate a small difference between placebo and analgesia treatment in self-reported pain unpleasantness (Fig. 2). This difference is in the expected direction and is explained by greater analgesia after the administration of the real analgesic (lidocaine) than the placebo treatment (vaseline). Interestingly within placebo responders, the treatment effect size (i.e. treatment vs. control) was larger for placebo than lidocaine for pain unpleasantness (d=-0.53 vs. −0.21). This unexpected pattern may have been driven by the greater pain unpleasantness rating in the placebo control condition, compared to the analgesic control condition (Fig. 2 B). This difference is unlikely to be explained by response bias and social desirability (Hrobjartsson et al., 2011), as participants were on the assumption that both creams were analgesics.

The current design allows us to parse the magnitude of placebo analgesia by not only comparing the inert cream against an actual analgesic but also accounting for the variability associated with the stimulation of mirror body territories which were not treated with the inert cream or actual analgesic (Fig. 1), in a sample (n=39) larger than the majority of similar previous studies. Our results indicate a small non-significant difference between placebo and the actual analgesic condition as reflected by ratings of pain unpleasantness of pain (Fig. 2). Interestingly, the control conditions revealed a trend similar to the treatment conditions (namely analgesia lower than placebo). This was accounted for by greater pain unpleasantness during the placebo-control condition compared to all the other conditions (Fig. 2 B). The N2-P2 LEPs confirmed that the most important factor explaining variability of these neural responses was the expectation of being treated with an analgesic cream, regardless of whether this cream was a real analgesic or just vaseline (Fig. 4).

These findings provide further evidence in support of the response expectancy theory (Kirsch, 1997; Koyama, McHaffie, Laurienti, & Coghill, 2005; Montgomery & Kirsch, 1997). Akin to other studies we provided our volunteers with positive expectation about the treatment and did not implement a conditioning procedure (De Pascalis, Chiaradia, & Carotenuto, 2002; Paul Enck, Bingel, Schedlowski, & Rief, 2013; Pollo et al., 2001). On the contrary, we implemented a well-established script of verbal suggestion within a ritual context (see appendix) that led the majority of healthy volunteers to believe in the experience of a reduction of pain following administration of an inert cream, particularly a decrease in the affective component of their sensation. Interestingly, we observed a greater difference between placebo treatment and control (namely, a greater reduction of pain) than between analgesic treatment and control (Table 1). Individuals who showed a greater self-reported placebo effect as measured with the pain unpleasantness ratings also demonstrated a greater modulation of the N2-P2 amplitude for placebo treatment (Fig. 4). This robust positive relationship between the behavioral and the neural marker provides an index of convergent validity for the reported results.

An alternative interpretation of the current results can also be based on a “nocebo” effect associated with the control (i.e. no treatment) conditions. Such an interpretation would suggest that individuals who experienced a lower placebo effect had greater negative expectation from the pain stimulation on the control limb, and this correlated with the extent of the N2-P2 modulation. Other authors have similarly speculated that the placebo and nocebo conditions may be used by experimental volunteers as reference perceptual criterion against which compare the sensations experienced during the “neutral” control condition (Freeman et al., 2015). Future studies may address not only the role of implicit and explicit positive expectations in triggering and maintaining placebo analgesia but also the role of co-occurring implicit contextual negative expectations that may arise from the stimulation of non-treated body parts. This observation leads us to two important caveats. First, the significance of these findings, and more generally of those obtained in the context of laboratory experiments on healthy volunteers, should not be generalized to the understanding of placebo responses in pain patients. In fact, a lack of correlation between placebo analgesia in experimental pain and clinical pain has been reported (Muller et al., 2016). Second, the interpretation of placebo effects is context-dependent and importantly relies on individuals’ interpretation of the treatment context (Enck & Klosterhalfen, 2013; Whalley, Hyland, & Kirsch, 2008). Consequently, different experimental designs can affect participant’s interpretation to a different extent and contribute to differences in the magnitude of the placebo effect.

Notwithstanding these caveats, our experimental design allowed us to precisely test the size of the placebo effect by calibrating it against a true analgesic. The experimental design allowed a head-to-head comparison between the analgesic and the placebo, due to the presence of both a real analgesic compound and of a non-treated skin territory on a body area exactly contralateral to the experimentally treated one. Unfortunately however, this design does not allow us to examine the earliest response to nociceptive stimuli, as measured through the N1 component (Valentini et al., 2012) as upper and lower limbs are associated with different arrival time in the somatosensory cortices, and thus with different latencies of the evoked brain signals. Hence we focused on the magnitude of the N2 and P2 potentials for the current study. It is noteworthy that the majority of previous studies report a reduction of the N2 and P2 potentials during placebo analgesia (Colloca et al., 2008; Martini, Lee, Valentini, & Iannetti, 2015; Wager, Matre, & Casey, 2006; Watson, El-Deredy, Vogt, & Jones, 2007).

In conclusion, our findings provide an ecologically valid estimate of the placebo analgesia effect by comparing a placebo treatment directly against that of a real analgesic. We show that verbal suggestions alone are sufficient to establish a moderate placebo effect and that unpleasantness of pain is the most sensitive measure of the placebo analgesia. We also show that the EEG measures of placebo analgesia are strongly correlated with the magnitude of the placebo analgesia computed from pain unpleasantness ratings. Future studies should examine individual differences in the behavioural and neural measures of placebo analgesia.

## Appendix

### Induction script

“Thanks for coming. You are volunteering for the final phase of a clinical evaluation of two new analgesics, *Exacaine* and *Varicaine* (these are the commercial labels and the active component cannot be disclosed). The active components are completely harmless and have no side effects in humans. You will participate in a study in which we will be testing the efficacy of a new analgesic technique on the experience of pain and on brain activity. During the experiment we will deliver thermal (laser) stimuli which can induce pricking and hot sensations. These sensations may be interpreted as painful depending on your very personal estimate. Importantly, we will use only one stimulus energy during the experiment, which will correspond to what you will judge as a moderate sensation of pain. We will spread one cream on one limb and the other cream on another limb. It will take about 10 minutes to come into action. Afterwards we will rub it off from your skin and start with the stimulation protocol”.

## Acknowledgments

We thank Prof. Simon Baron-Cohen, Dr. Karthik Bhargavan, Dr Giuseppina Porciello, Ms Assunta Ruggiero, Dr. Li Hu, for their help and encouragement at various stages of this project. E Valentini and SM. Aglioti were supported by Italian Ministry of University and Research (PRIN, Progetti di Ricerca di Rilevante Interesse Nazionale, 2015, Prot. 20159CZFJK). B Chakrabarti was supported by a British Council Researcher Exchange grant.

